# Paper-Based Microfluidic Continuous-Flow Electroporation (µCEP) for Bacterial Transformation through Pore-Scale Electric-Field Enhancement

**DOI:** 10.64898/2026.10.19.760286

**Authors:** Bhavya Jaiswal, Md. Nazibul Islam, Zachary R. Gagnon

## Abstract

Electroporation (EP) is an established method for introducing biomolecules such as DNA into bacterial cells. Commercially available cuvette-based EP systems are expensive to use and limited to small sample volumes making them impractical for high-throughput applications. Furthermore, high electric fields are required in these systems which can promote electrolysis and Joule heating, thereby reducing cell viability. Herein, we report a scalable, low-cost technique called Microfluidic Continuous-Flow Electroporation in Paper (µCEP) which takes advantage of the insulating nature of glass fiber paper and its micron-sized pores to amplify the electric field strength experienced by cells. The µCEP device is fabricated by sealing glass fibers between PDMS membranes using a combination of corona plasma treatment and heat-pressing. Our results demonstrate that the µCEP technique enables transformation of *E. coli* at lower applied electric field strengths with transformation efficiencies comparable to or higher than those of cuvette-based EP under the tested conditions. The continuous-flow format and lower required applied field provides a low-cost approach to bacterial electroporation while maintaining cell viability comparable to that of conventional cuvette-based EP.

## Introduction

Genetic engineering of organisms has a variety of applications such as microbial therapeutics (1,2), cellular reprogramming (3,4), sustainable agriculture (5,6), biofuel production (7,8), and biomass production (6,9). Electroporation (EP) is a widely used method for genetic engineering that applies a quick high-voltage pulse (typically over 1 kV) across electrodes separated by a millimeter-scale gap, thus generating a strong electric field (10). This field increases the local transmembrane voltage (TMV), disrupting the cell membrane; when the TMV exceeds the critical transmembrane potential, hydrophilic pores form and increase membrane permeability, allowing foreign material to enter the cell. Under optimized conditions the pores reseal, yielding viable transformed cells (11).

Conventional cuvette-based EP, first reported in 1982 (12,13), is simple to use and widely available, yet its format remains largely unchanged and suffers from several drawbacks. High-voltage pulses cause Joule heating and electrolysis, which damages the cell membrane and creates bubbles, respectively (14). Moreover, the electrolysis-derived ions and degradation of aluminum electrodes is toxic and shifts the pH, thus decreasing cell viability (15). Moreover, since cuvettes can only accommodate small sample volumes (∼20–100 µL), high-throughput workflows become costly and impractical; generating a mutant library from a few milliliters of cell suspension would require hundreds of individual cuvettes. This stagnation contrasts sharply with the rest of the genetic engineering toolkit, where CRISPR screening (16), next-generation sequencing (17,18), and microfluidic cell-handling platforms (19,20), have all scaled to process thousands to millions of samples in parallel. As these upstream and downstream technologies have advanced, cuvette-based EP has increasingly become the bottleneck in end-to-end workflows, constraining throughput at the cell-engineering step.

The recent expansion of microfluidic technology has renewed interest in EP platform development. Microfluidic EP (µEP) platforms address the limitations of cuvette-based EP systems by confining the electric field to the scale of individual cells (10,21–25). This achieves the required field strength at substantially lower voltages, which reduces Joule heating and electrolysis, thereby improving cell viability (10). A common approach is to use MEMS (micro-electro-mechanical systems) fabrication to shrink the interelectrode spacing to the microscale (10,21–25) or to build structures such as micronozzle/trap arrays (26,27) and nano-spikes (28). Yet, MEMS fabrication is laborious, costly, and difficult to scale, posing a technological and financial barrier to commercialization (22,29). To avoid MEMS fabrication, Cao et al. developed a low-cost platform in which cells on a commercial track-etched water filter membrane are placed between flat electrodes, focusing the electric field through the nanopores to achieve effective localized delivery (30). Nevertheless, the small single-well format limits throughput, a limitation shared by many µEP platforms (10,22,24). Scaling µEP to high-throughput operation requires careful control of microchannel and electrode geometries, as these parameters influence electric-field strength and spatial distribution and, consequently, the conditions experienced by cells during continuous flow (22,25). Recently, a simple µEP device capable of high-throughput operation was reported in which plastic tubing between two syringe needles matched or outperformed cuvette-based EP across several conditions (31). However, in this design the field is set solely by the applied voltage and electrode gap, with no reported geometric focusing of the electric field. There remains a need for low-cost, high-throughput µEP platforms that concentrate the field at the cellular scale, achieving the field strengths required for efficient bacterial electroporation at substantially lower applied voltages, which may reduce Joule heating and electrolysis while maintaining or improving transformation efficiency and cell viability.

Over the past decade, microfluidic technology has expanded towards the development of disposable, ultra-low-cost paper-based devices (32,33). To address the cost, throughput, and field-focusing limitations of cuvette-based EP and existing µEP platforms, we developed Microfluidic Continuous-Flow Electroporation in Paper (µCEP), a paper-based microfluidic platform for continuous EP. This is based on the microfluidic pressure in paper (µPiP) technology developed by our lab, which uses encapsulated paper fluidic channels that can withstand continuous pressure-driven fluid flow, enabling continuous-flow operation (34–37). Instead of relying on MEMS fabrication, the µCEP platform takes advantage of the insulating nature of glass fiber paper, whose inherent porous structure focuses electric fields without requiring any additional treatment. The low-cost paper consists of a non-woven network of compressed glass fibers, which forms numerous micron-sized pores that amplify the local electric field, enabling effective EP of bacterial cells at low applied voltages. Our device is ultra-low-cost (30 ¢ per device), processes larger sample volumes than conventional cuvettes, and can be easily parallelized for high-throughput EP. To the best of our knowledge, this is the first instance where paper microfluidic devices have been used to enhance electric field strengths for EP. We believe µCEP will help democratize EP, making it accessible to educational institutions and academic labs in low-resource settings. We first use micro-computed tomography (micro-CT) and finite element analysis (FEA) to identify regions of enhanced electric field within the micron-scale pores of the glass-fiber substrate. As a proof-of-concept, we investigated the transformation efficiency of the µCEP platforms for DH5α cells under various electric field strengths. Under the tested conditions, µCEP reduced the projected consumable cost for processing 1 mL from $62.50 to ∼$0.30 (∼208-fold, calculations provided in Supplementary Table S1 and S2), achieved >5-fold higher transformation efficiency at selected matched field strengths, and reduced the minimum applied field required for quantifiable transformation from 6 to 2 kV/cm (3-fold). Finally, we demonstrate the application of the µCEP platform on *E. coli* K-12 wildtype which is more challenging to engineer and observed similar results (38).

## Materials and Methods

### Micro-CT Analysis of Paper Channels

The glass fiber substrate was quantitatively imaged using micro-CT to study the porous structure of the insulating fibers and the resulting electric field distribution. Scans were carried out using the SkyScan 1272 bench top micro-CT scan (Micro Photonics Inc.; 1.5 µm voxel size, 800 ms exposure time) and CTvox software was used for image processing. The resulting micrographs were digitally converted into a 2D mesh and used as a model for FEA using COMSOL Multiphysics software v6.1 (COMSOL, Burlington MA, USA). Each computational model consisted of a 2D slice with an element mesh consisting of 747109 triangular and 69550 quadrilateral mesh elements.

### Electric Field Distribution and Velocity Field Analysis

Both the bulk fluid flow and applied electric field were oriented along the longitudinal axis of the paper channel, since the steel tubes served as inlets/outlets for the cell suspension and were also used as electrodes. Fiber surfaces were set as electrical insulators and water was assigned within the bounded domain. Under the assumptions of a homogeneous and isotropic medium with no internal current or charge sources, the distribution of the electric potential simplifies directly to Laplace’s equation:

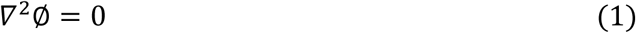

Where ∅ is the electric potential and the electric field was determined by:

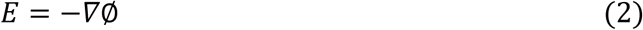

One side of the paper structure was defined as the ground and boundary potentials corresponding to applied electric field of 6 kV/cm was imposed across the modeled domain. A field strength of 6 kV/cm was chosen as a conservative lower bound, consistent with the experimentally observed threshold voltage for EP (see Fig 4a and 5a) and is within the range of fields used in previous *E. coli* EP studies (39,40). The resulting electric field distribution was used to investigate the electric field enhancement within the glass fiber substrate.

To characterize fluid transport through the porous glass-fiber structure, a steady-state creeping-flow simulation was performed using the 2D micro-CT-derived geometry. The fluid was modeled as incompressible and Newtonian, with a density of 1000 kg/m³ and a dynamic viscosity of 1 × 10⁻³ Pa·s (41). Since the characteristic Reynolds number was much smaller than unity, fluid inertia was neglected, and the flow was described using the Stokes momentum and continuity equations:

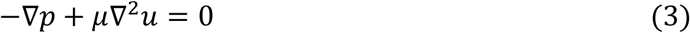

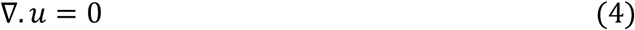

Where *p* is fluid pressure, *μ* is dynamic viscosity, and *u* is the fluid velocity vector. For the modeled boundary conditions, a uniform normal inlet velocity of 5.68 × 10⁻⁴ m/s was prescribed, the outlet was assigned zero relative pressure, and the glass-fiber surfaces were treated as stationary no-slip boundaries. The resulting velocity distribution was used to identify flow-accessible regions for subsequent analysis of the local electric-field distribution.

### Device Fabrication

The fabrication of the µCEP device takes less than 10 minutes, and is based on a previously developed fabrication technique called microfluidic pressure in paper (µPiP) (Fig 1A) (35). First, square PDMS sheets with 0.1 mm holes and elliptical microfluidic paper channels (3mm wide, 5 mm long) were cut from sheets of electronic grade fiberglass Craneglass® 230 paper (Neenah Filtration) using a CO_2_ laser cutter (LS-2440, Boss Laser). Next, the glass fiber channel (Fig 1B) was aligned and laminated between the two polydimethylsiloxane (PDMS) sheets (0.5 mm, McMaster-Carr). The two sheets were oxidized and irreversibly bonded to each other using the oxygen plasma generated by a handheld Tesla coil (Electro-Technic Products, Model BD-20AC). Finally, to remove any air gaps between the PDMS sheets, the device was placed between a bench top heat press (Dulytek DW 400) at a temperature of 55°C for 5 minutes. A circular PDMS piece with two 0.1 mm holes created using a biopsy punch (Ted Pella, Inc) is irreversibly bonded to the inlet and outlet using a Tesla coil as shown in Fig 1C. This enables the inlet and outlet to be connected to a small 1 cm long, 0.064 mm ID stainless steel tube (New England Small Tube). A 2D CAD drawing of the device with the dimensions of all components is provided in Supplementary Fig S1. The µCEP device was placed in a petri dish and sterilized in a biosafety cabinet with UV irradiation overnight before use. For experiments, EP pulses were delivered to the tube while a syringe containing the cell suspension was connected to the tube using a 0.1 mm ID tubing (Cole Palmer) to introduce the cells.

**Figure 1.**
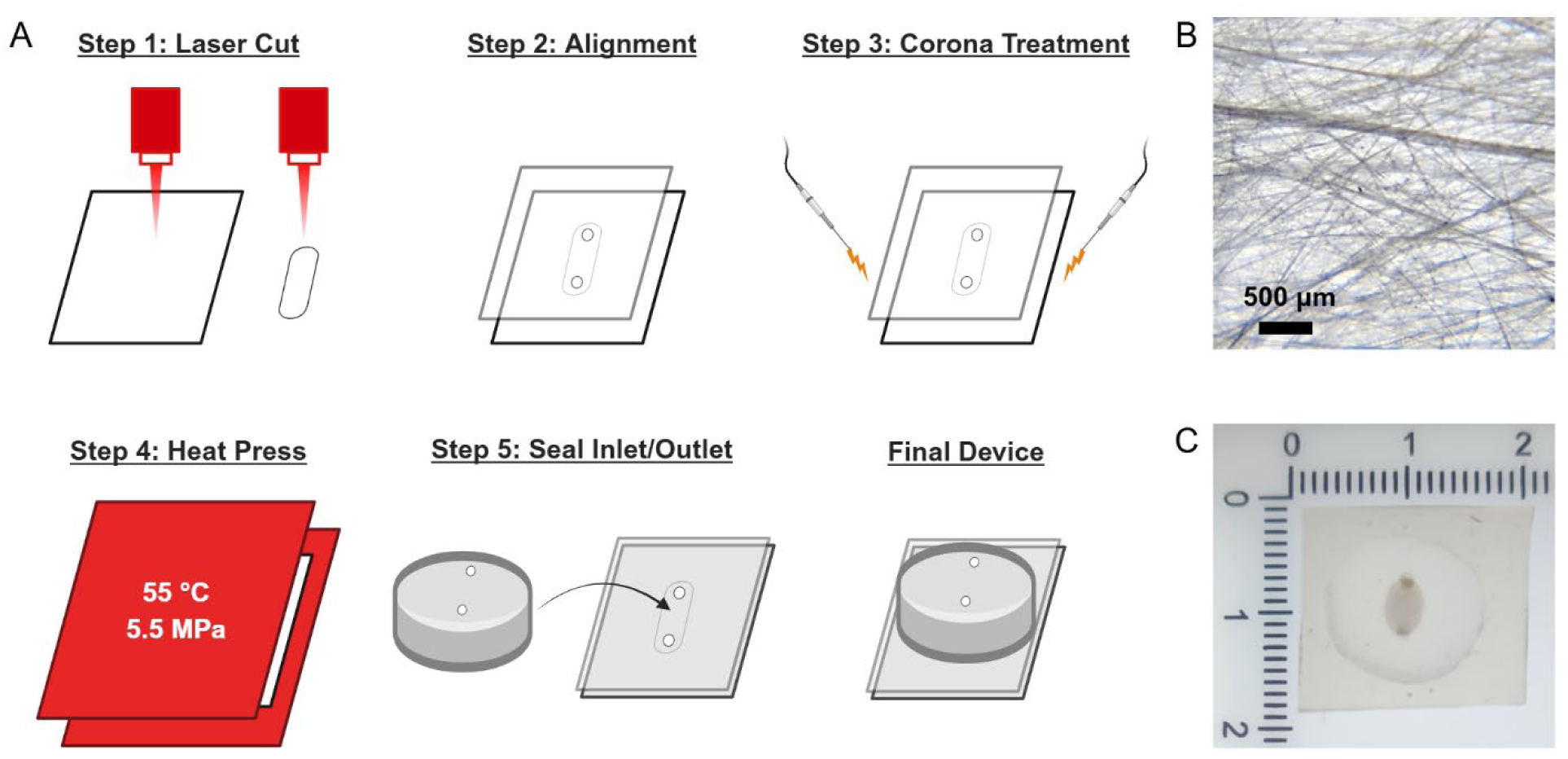
Schematic of the Microfluidic Continuous-Flow Electroporation in Paper (µCEP) device. A) Fabrication workflow using the µPiP technique. B) Glass fiber substrate imaged using bright-field microscopy. C) Completely assembled and sealed µCEP device.

### Cell Culture and Preparation for Electroporation

Experiments were performed with two *E. coli* strains DH5α (Thermo Fisher Scientific, MA, USA) and K-12 strain MG1655 (*E. coli* Genetic Stock Center, Yale University), which were made electrocompetent via the standard methods (42). Briefly, bacterial cultures were grown overnight in 50 mL LB media at 37°C with shaking. The next day, 25 mL of the overnight culture was transferred to 500 mL LB media and incubated at 37°C with shaking to exponential phase (OD_600_ ∼ 0.4). Cultures were chilled on ice and centrifuged at 1000g and 4°C for 15 min using a Heraeus^TM^ Multifuge^TM^ X1 centrifuge with a TX-400 rotor (Thermo Scientific, Rockford, IL, USA). The supernatant was discarded, and the cells were resuspended in ice-cold Milli-Q water. The cells were then washed three times in 10% glycerol under the same conditions. The final suspension was diluted to an OD_600_ of ∼100 in 10% glycerol and was stored at −80°C until use. Cell concentration was confirmed by measuring the OD_600_ at a 1:100 dilution using a NanoDrop One Microvolume UV–Vis Spectrophotometer (Thermo Fisher Scientific, MA, USA). Viable cells were quantified on LB agar, and transformants were quantified on LB agar containing 100 µg/mL ampicillin. pUC19 plasmid (Thermo Fisher Scientific, MA, USA) was diluted to the appropriate concentration in 1× Tris-EDTA (TE) buffer (10 mM Tris-Cl, pH 8.0; 1 mM EDTA).

### Cell Electroporation

For electroporation, cells were thawed on ice and diluted 20× in 10% glycerol. pUC19 was added to the cell suspension to a final concentration of 100 pg/µL and mixed gently by pipetting. Using a pre-chilled 1 mL syringe (BD), 200 µL of the cell suspension was flowed through a chilled µCEP device at 50 µL/min with a syringe pump (Chemyx Fusion 100). EP pulses were delivered through the inlet and outlet steel tubes using a waveform generator (RIGOL DG4102). Square wave signals from the function generator were amplified 200-fold by a high-voltage amplifier (TREK Model 677B high-voltage power amplifier, Trek Inc., Lockport, NY) and monitored using an oscilloscope (RIGOL DS1054) (Fig 2). EP pulses had a time constant of 6 ms, time period 20 ms, and lead and trailing times of 40 µS. The function generator’s maximum output limited µCEP to field strengths up of ≤6 kV/cm. Cells at the outlet were collected in a centrifuge tube containing pre-warmed (37°C) LB media. A further 100 µL of LB was flowed through the µCEP device after EP to collect any remaining cells. The recovered suspension was transferred to a culture tube and incubated for an hour at 37°C. Cells were then plated on LB agar plates and incubated overnight.

**Figure 2.**
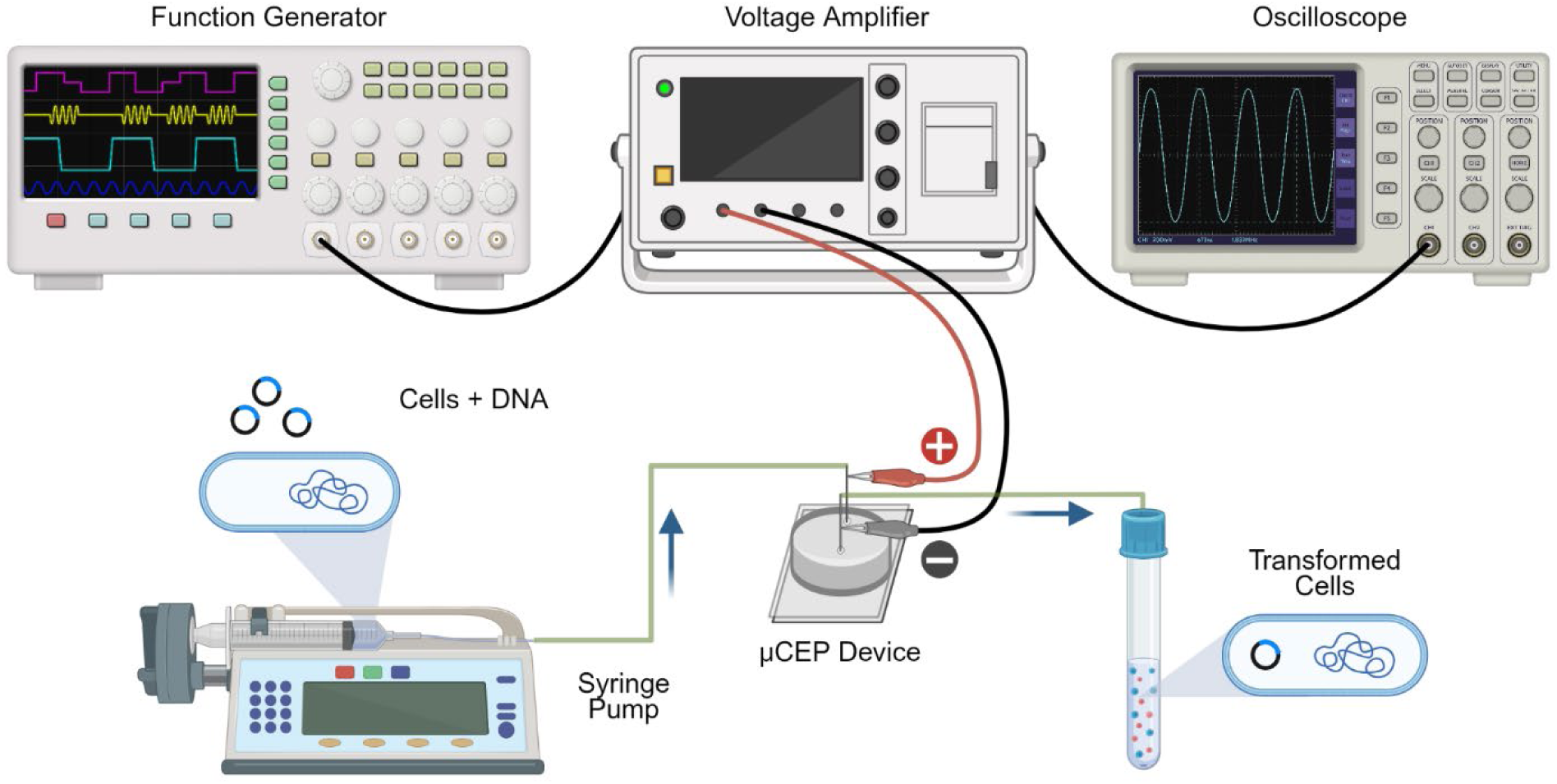
Schematic of the µCEP setup. A programmable syringe pump drives the cell suspension through the µCEP device and into the recovery media. A function generator produces continuous electrical signals which are amplified by a high-voltage power amplifier and delivered to the µCEP device.

As a positive control, 20 µL of the same cell suspension containing DNA was pipetted into pre-chilled 0.1 cm EP cuvettes and electroporated at electric field strengths between 2–8 kV/cm using a Nepa Gene ELEPO21 electroporator (Bulldog Bio, Portsmouth, NH). Immediately after the electric pulse was delivered, prewarmed (37°C) LB media was added to the cuvette and mixed with the cells via pipetting. The recovered suspension was transferred to a culture tube and incubated and plated according to the protocol mentioned earlier.

Images of agar plates were quantitively analyzed with the assistance of open-source software OpenCFU (43). Transformation efficiency was calculated for each technical replicate as the number of colony forming units (CFU) divided by amount of DNA plated (in µg) and reported as CFU/µg-DNA. Cell viability was determined by normalizing the total CFU count for each condition to that of the no pulse control.

### Statistical tests

Transformation-efficiency (CFU/µg-DNA) values were log10-transformed for statistical testing, as efficiencies vary multiplicatively. µCEP and cuvette-based EP were compared using a two-sided Welch’s t-test on triplicate measurements.

## Results

### Finite Element Analysis of Electric Field Strength Enhancement in Glass Fiber

First, to investigate how the porous structure of the paper affects the local electric field strength, we used FEA on a digitized geometry of the glass fiber substrate. A micro-CT scan of the paper was reconstructed into a 3D image, and a 2D slice of this scan was used as the computational domain as shown in Fig 3A.

**Figure 3.**
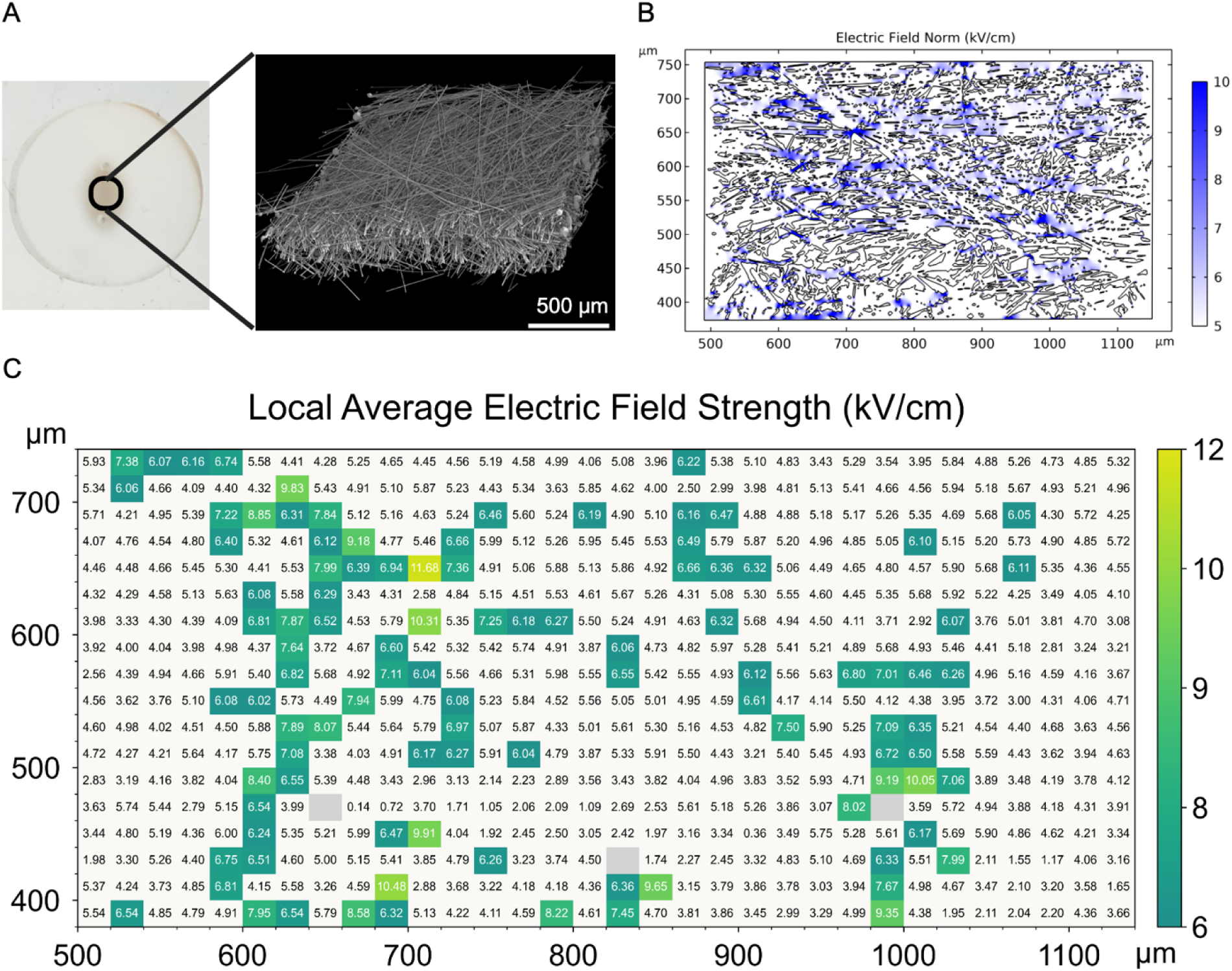
FEA of the electric field enhancement within the glass fiber substrate. a) Micro-CT reconstruction of the porous glass-fiber structure (voxel size: 1.5 µm). b) Simulated electric field magnitude within the micro-CT derived 2D geometry at a nominal applied field of 6 kV/cm, illustrating localized field enhancement near insulating fiber boundaries. Plot is re-scaled to highlight areas of high electric field strength. c) Spatial distribution of bin-averaged electric field magnitude in regions satisfying the flow-velocity criterion (>30 µm/s). Field was averaged over 20 × 20 µm bins to quantify local variation relative to the applied field of 6 kV/cm. Colored bins have an average field above the applied field, with color indicating magnitude; white bins are at or below it. Gray bins had insufficient data (>95% of data points outside the selection criteria).

Solving Eqns. 1 & 2 on this geometry revealed localized field enhancement near narrow gaps and sharp fiber features, where the insulating geometry concentrates the field relative to the nominal applied field (Fig 3B and Supplementary Fig S2). The color scale in Fig 3B was rescaled to highlight regions where the local field exceeds the applied 6 kV/cm. This is consistent with earlier reports that in insulating structures with sharp ends there is a swift change in the radius of curvature (ROC) which constricts the electric field (44). This produces stronger local electric field strength as compared to rounded ends where the ROC changes more gradually. Since the field experienced by a cell depends on the path it takes through the paper, next we characterized the fluid flow. Although *E. coli* swim at approximately 20–22 µm/s, their transport through confined structures is significantly influenced by flow conditions (45,46), so the flow field is needed to predict likely transport pathways.

Solving Eqns. 3 & 4 revealed several zones of high velocity that form preferential flow paths through the paper (Supplementary Fig. S3). In confined porous geometries, bacteria tend to accumulate and remain trapped in regions where local flow velocities are comparable to or below their swimming speed (47). Below this threshold (≤10 µm/s), bacteria exhibit strong hydrodynamic trapping near surfaces, while above ∼40 µm/s, shear forces overcome these trapping mechanisms and bacteria are dislodged into the bulk flow (47). To focus on regions where cells are likely to be mobile rather than trapped, we restricted the electric field analysis to locations where the velocity exceeded 30 µm/s and averaged the field in 20 µm × 20 µm bins. The bin size corresponds to the distance an *E. coli* cell advected at 4000 µm/s travels during a 5 ms EP pulse. Of the filtered data points, 99.8% had fluid velocities below this value (the remaining 0.2% did not exceed 7000 µm/s), thus cells in nearly all flow-accessible regions are displaced by less than one bin width during an EP pulse.

Based on these criteria, we calculated the average field strength in each bin, excluding regions along the edges of the computational domain to avoid artifacts from boundary effects. (Fig 3C and Supplementary Fig S4). The color scale in Fig 3C was rescaled to highlight regions where the local field exceeds the applied 6 kV/cm. Three bins (gray in Fig 3C) were excluded because more than 95% of their data points fell outside the velocity criteria resulting in insufficient data. Among the remaining bins, the field was heterogenous, with 17.5% of bins exceeding the applied 6 kV/cm, with a maximum bin-averaged value of 11.68 kV/cm, while the rest of the bins fell below the applied field (white bins in Fig 3C). Since a substantial fraction of the flow-accessible regions experience fields above the applied value, cells passing through these regions may reach the field required for EP at a lower applied voltage than the nominal field suggests. Reducing the required applied voltage could in turn limit Joule heating and pH changes associated with pulsed electric fields, which can reduce cell viability and transformation efficiency, although these effects were not modeled here.

We note that this approach uses a 2D slice of a 3D structure, which can alter pore connectivity thereby influencing both flow paths and field magnitudes. The velocity threshold and binning serve as a proxy for the region cells are likely to traverse and particle tracing would give a more direct estimate of the field experienced along individual trajectories. Without setting the velocity threshold, the same analysis yielded 9.4% of bins exceeding the applied 6 kV/cm, while the rest of the bins fell below the applied field (Supplementary Fig S5).

### Performance of the µCEP Device for Bacterial Electroporation

To establish the utility of µCEP, we chose *E. coli* DH5α for transformation, as it is well-known for its robust transformation efficiency. First, we were curious whether *E. coli* could be mechanically transformed by the shear stress imposed by the micron-sized pores of the glass fibers. However, flowing cells through the µCEP device at fluid flow velocities up to 800 µL/min in the absence of an electric field did not result in any transformation events. We then evaluated how the µCEP device performed across a range of electric field strengths and observed that, at electric field strengths greater than 2 kV/cm, colonies were obtained with transformation efficiencies ranging from 10^6^ to 10^7^ colony-forming units (CFUs)/µg of DNA. Plates containing fewer than 30 colonies were considered below the quantification threshold and are reported as “too few to count”.

µCEP yielded higher transformation efficiency than cuvette-based EP at 6 kV/cm and quantifiable transformation at lower electric field strengths (Fig 4A). Cuvette-based EP produced quantifiable transformation only at field strengths ≥6 kV/cm, with transformants below the quantification threshold at ≤5 kV/cm, whereas µCEP yielded quantifiable transformation across the range 2-6 kV/cm. Since µCEP could not be tested above 6 kV/cm owing to the output limit of the function generator, 6 kV/cm was the only field strength at which the two methods could be compared directly. At 6 kV/cm, Welch’s t-test gives p-values of 0.0195 which indicates that µCEP showed significantly higher transformation efficiency than cuvette-based EP. Notably, µCEP at 2 kV/cm achieved efficiency similar to cuvette-based EP at 6 kV/cm thereby reducing the minimum applied field required for quantifiable transformation of DH5α by at least 3-fold, from 6 to 2 kV/cm.

**Figure 4.**
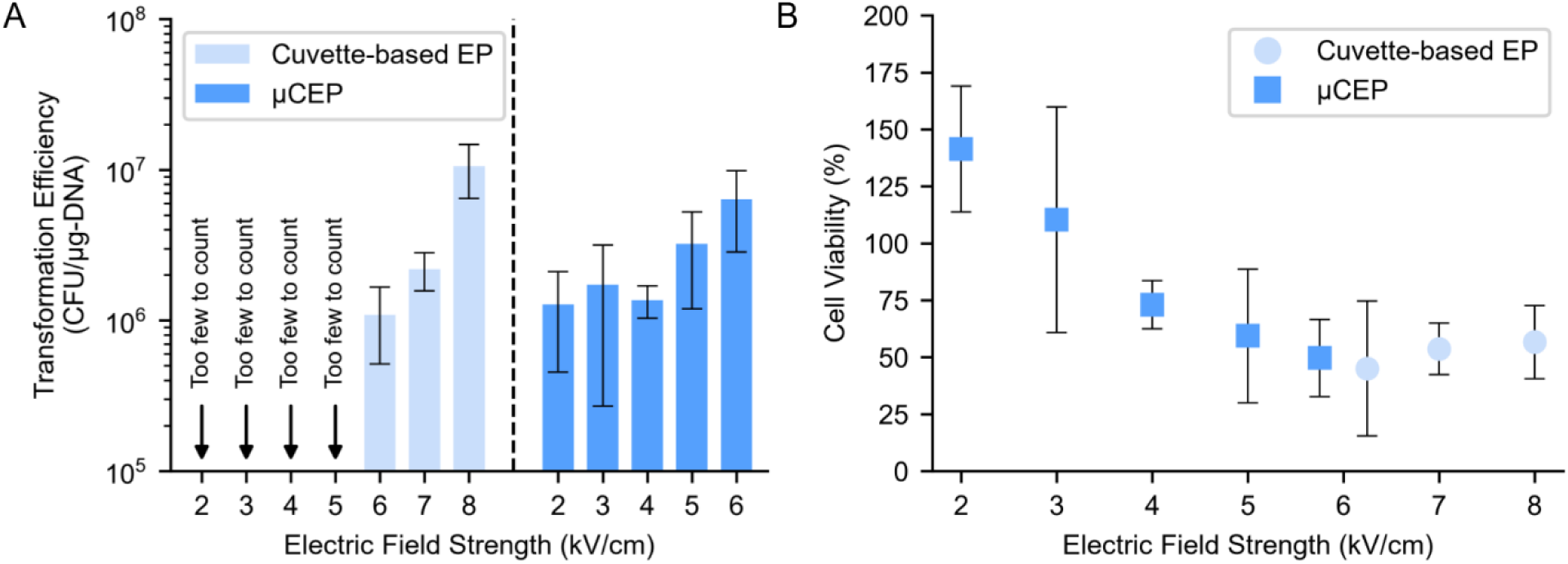
A) Transformation efficiency and B) cell viability after electroporation of pUC19 vector into *E. coli* DH5α via the µCEP device and cuvette-based electroporation (n=3). Plates containing fewer than 30 colonies were considered below the quantification threshold and are reported as “too few to count” (TFTC). n denotes the number of technical replicates; error bars indicate standard deviation.

Over the range of 4-8 kV/cm, cell viability after EP was of similar magnitude for the µCEP device and for cuvettes (Fig 4B). We hypothesize that the improved transformation efficiency of µCEP relative to cuvette-based EP is due to the synergistic effect of the continuous flow, which carries cells away from the electrodes, combined with the continuous application of electrical pulses. At applied field strengths ≤3 kV/cm, relative viable-cell recovery exceeded the no-pulse control, reaching approximately 145% at 2 kV/cm for DH5α. As this metric is based on CFUs measured after a 1-h recovery period, values >100% indicate recovery of more colony-forming cells than in the corresponding no-pulse control and should not be interpreted as survival exceeding 100%. Previous studies have shown that electrical stimulation can alter bacterial proliferation, although under conditions that differ substantially from ours. Brief localized stimulation selectively increased proliferation of motile *Bacillus subtilis* cells in biofilms (48). Additionally, sustained current densities of up to 10 A m⁻² A accelerated *E. coli* growth (shortening both lag period by up to ∼40% and doubling time by up to ∼70%) whereas densities above 20 A m⁻² impaired growth (49). Therefore, the increased CFU recovery observed at low field strength may reflect differential post-pulse growth, or a stimulatory response to electrical exposure; distinguishing among these possibilities will require time-resolved measurements of cell number after EP. The proliferation of bacteria in response to electrical stimulation is beyond the scope of the current work; however, identifying the conditions that enable this phenomenon would be interesting to investigate in the future. We also tested different pretreatment methods for the glass fiber paper by incubating it in solutions containing BSA in various concentrations between 0-5% w/v. Interestingly, BSA did not have any significant impact on transformation efficiency (data not shown).

To determine whether the low-field performance of µCEP extended beyond DH5α, we next tested the K-12 strain *E. coli* MG1655, which is less readily transformed than DH5α, since it contains native restriction and modification systems for DNA methylation and degradation, thereby making it more challenging to engineer (38). First, we evaluated how the µCEP device performed across a range of electric field strengths and observed that, at electric field strengths greater than 2 kV/cm, colonies were obtained with transformation efficiencies ranging from 10^5^ to 10^7^ colony-forming units (CFUs)/µg of DNA.

Similar to the performance observed with the DH5α strain, µCEP yielded higher transformation efficiency than cuvette-based EP at 6kV/cm and quantifiable transformation at lower electric field strengths (Fig 5A). Cuvette-based EP produced quantifiable transformation only at field strengths ≥6 kV/cm, with transformants below the quantification threshold at ≤5 kV/cm, whereas µCEP yielded quantifiable transformation across the range 2-6 kV/cm. At 6 kV/cm, Welch’s t-test gives p-values of 0.0009 which indicates that µCEP showed significantly higher transformation efficiency than cuvette-based EP. Notably, µCEP at 2 kV/cm once again achieved efficiency similar to cuvette-based EP at 6 kV/cm, thereby reducing the minimum applied field required for quantifiable transformation of MG1655 by at least 3-fold, from 6 to 2 kV/cm. Relative viable-cell recovery decreased with increasing field strength for both methods; µCEP recovery decreased from approximately 135% at 2 kV/cm to approximately 30% at 6 kV/cm Fig 5B). Together, these results demonstrate that the reduced-field operation of µCEP is retained in another *E. coli* strain with lower baseline transformability.

**Figure 5.**
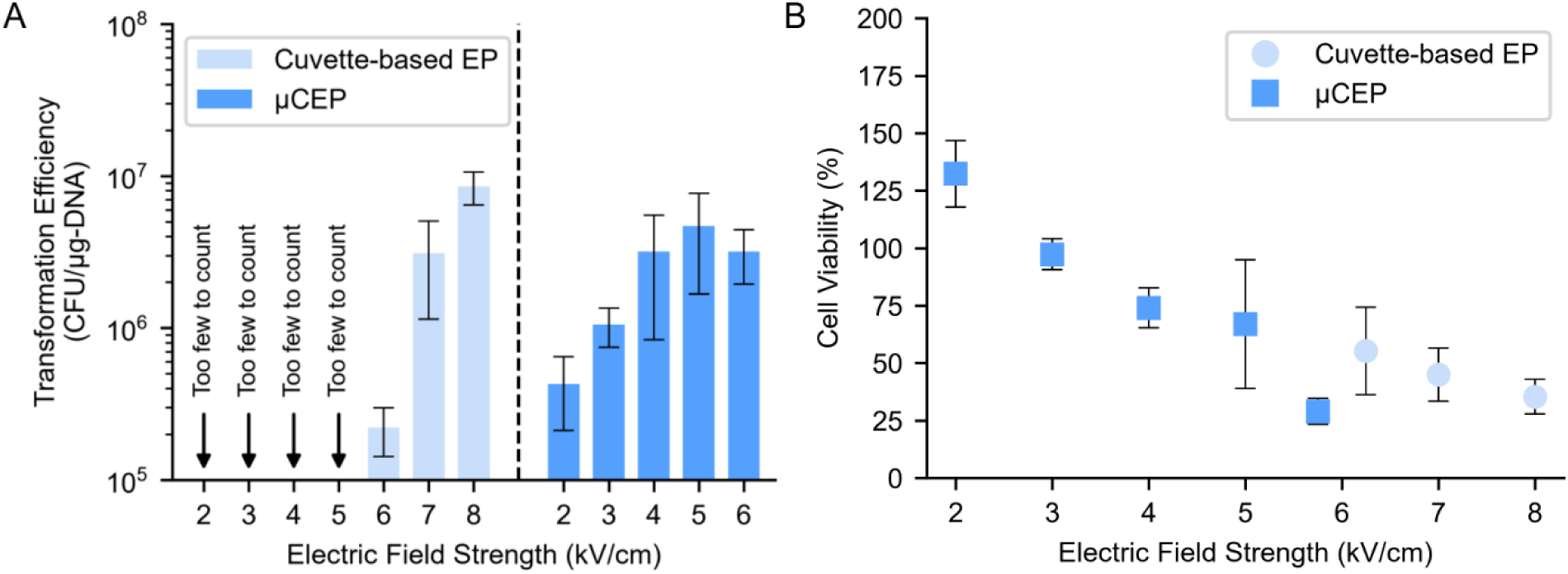
A) Transformation efficiency and B) cell viability after electroporation of pUC19 vector into *E. coli* K-12 MG1655 via the µCEP device and cuvette-based electroporation (n=3). Plates containing fewer than 30 colonies were considered below the quantification threshold and are reported as “too few to count” (TFTC). n denotes the number of technical replicates; error bars indicate standard deviation.

## Conclusions

Genetic engineering is a key process in several fields of scientific research, including cancer treatment, human microbiomes studies and pharmaceuticals. Improving the accessibility of cell transformation techniques can expedite researchers’ efforts in addressing various global challenges. The µCEP platform is ideal for cell transformation since it can be easily fabricated without complicated pretreatment steps, doesn’t require MEMS fabrication, and can be readily scaled up. It is an ultra-low-cost device that is well suited for bacterial EP and has the potential to extend the accessibility of EP to scientific research in low-resource settings or incorporated into university curricula for educational purposes.

We first examined how the microstructure of glass fiber paper affects the local electric field, using micro-CT scans of the substrate. FEA of a micro-CT-derived 2D geometry revealed that the insulating fibers concentrate the field in narrow pore gaps and near sharp fiber features, with 17.5% of the flow-accessible 20 µm x 20 µm bins exceeding the applied 6 kV/cm. Motivated by these findings, we developed Microfluidic Continuous-Flow Electroporation in Paper (µCEP), a device incorporating these glass fiber substrates, building on the µPiP technology previously developed by our group. In both *E. coli* DH5α and MG1655, µCEP yielded quantifiable transformation at applied fields as low as 2 kV/cm, whereas cuvette-based EP produced transformants below the quantification threshold at ≤5 kV/cm. At 6 kV/cm, the only field strength at which both systems could be compared directly, µCEP showed significantly higher transformation efficiency than cuvette-based EP and relative viable-cell recovery was comparable between the two systems. The ability to transform at lower applied fields is consistent with the simulated field concentration by the insulating glass fibers. Furthermore, the µCEP platform offers a higher throughput than commercially available cuvette-based EP, while being significantly cheaper.

Previously our lab demonstrated the use of low-cost paper devices for trapping bacteria via dielectrophoresis (DEP) (36). In future work, we plan to upgrade the µCEP platform by investigating the preconcentration of bacteria using DEP followed by EP. Additionally, it would be interesting to explore the potential for multiplexed EP to create mutant libraries (50–54) or for rapidly screening pulse waveforms to identify conditions that efficiently transform recalcitrant, non-model organisms. Eventually, we envision the development of an integrated, disposable lab-on-a-chip cassette capable of performing a range of synthetic biology techniques.

## Acknowledgements

The PI gratefully acknowledges support from a NASA Early Career Faculty award (80NSSC19K1401) and NSF I-Corps award (2112224).

## Author Contribution

B.J. and Z.G. conceived and designed the experiments. B.J. and N.I. carried out COMOL Simulations and analyzed the data. B.J. developed the µCEP platform, performed experiments and analyzed the data. B.J. and N.I. wrote the manuscript with input from Z.G.

