## Supplemental Data 1 for "Paper-Based Microfluidic Continuous-Flow Electroporation (µCEP) for Bacterial Transformation through Pore-Scale Electric-Field Enhancement"

**Table S1** Cost price of the parts of the µCEP device and conventional cuvettes

| **Device** | **Item** | **Price** |
| --- | --- | --- |
| Cuvette-based Electroporation | Electroporation chamber | $1,095 |
|  | 0.1 cm cuvette (Pack of 500) | $1.25* |
| Microfluidic Continuous  Electroporation | Constant Pressure system | ~$500 |
|  | µCEP Device | ~$0.30 |

*Lowest cost price of the cuvette is $1.56, a Price: Sales ratio of 0.8 has been assumed.

The electroporation pulse-generation and amplification hardware was excluded from the cost comparison because the operating principle behind the electrical systems for both cuvette-based EP and µCEP is the same.

Table S2 Cost to electroporate a unit volume sample

| **Processing Volume** | **Cuvette** | **µCEP Device** |
| --- | --- | --- |
| 1 mL | $62.50* | ~$0.30 |
| 5 mL | $312.50* | ~$0.30 |

*Lowest cost price of the cuvette is $1.56, a Price: Sales ratio of 0.8 has been assumed

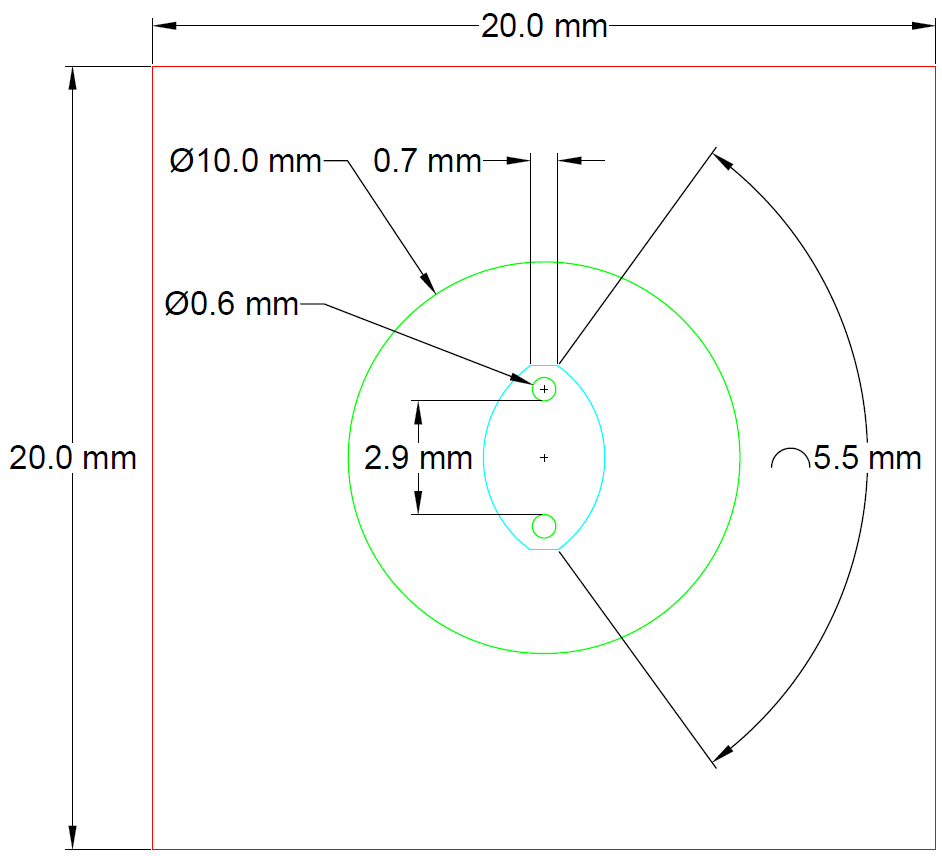

**Figure S1** CAD Drawing of the µCEP device. Red lines indicate the square PDMS sheet, green circle indicates the circular PDMS piece with holes for the electrodes and the cyan lines indicate the elliptical paper channel.

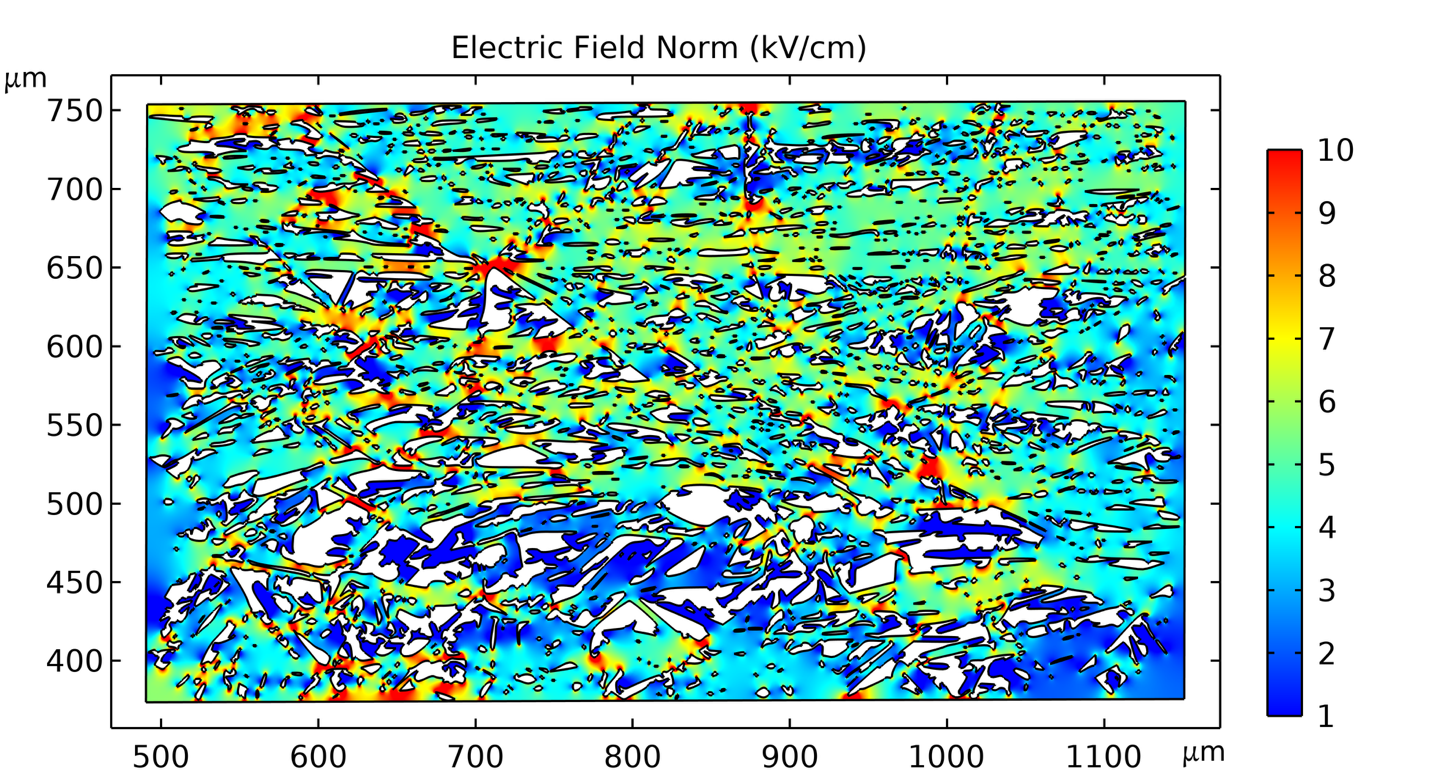

**Figure S2** Simulated electric field magnitude within the micro-CT derived 2D geometry at a nominal applied field of 6 kV/cm.

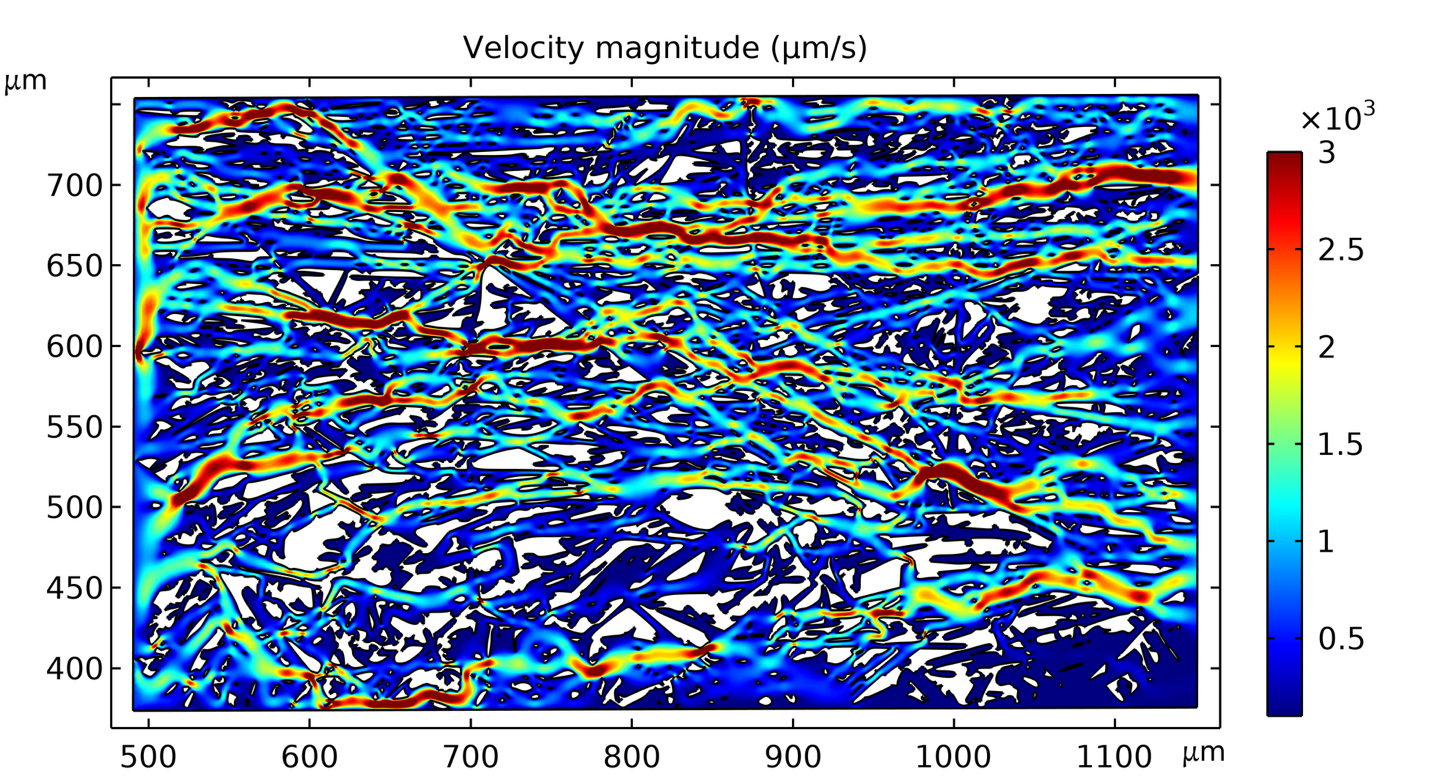

**Figure S3** Simulated velocity magnitude within the micro-CT derived 2D geometry computed for steady-state creeping flow.

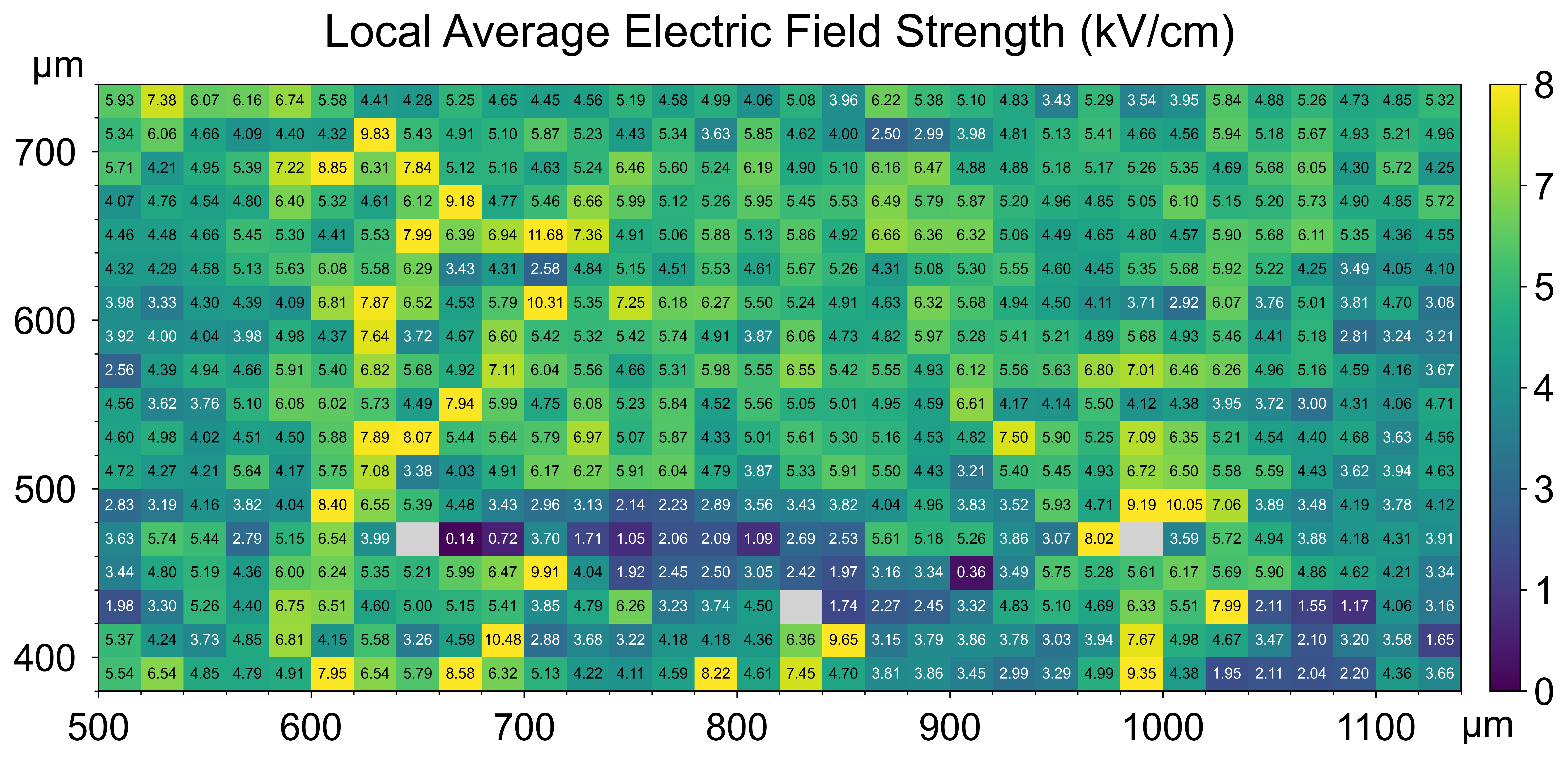
**Figure S4** Spatial distribution of bin-averaged electric field magnitude in regions satisfying the flow-velocity criterion (>30 µm/s). Field was averaged over 20 × 20 µm bins. Colored bins have an average field above the applied field, with color indicating magnitude. Gray bins had insufficient data (>95% of data points outside the selection criteria).

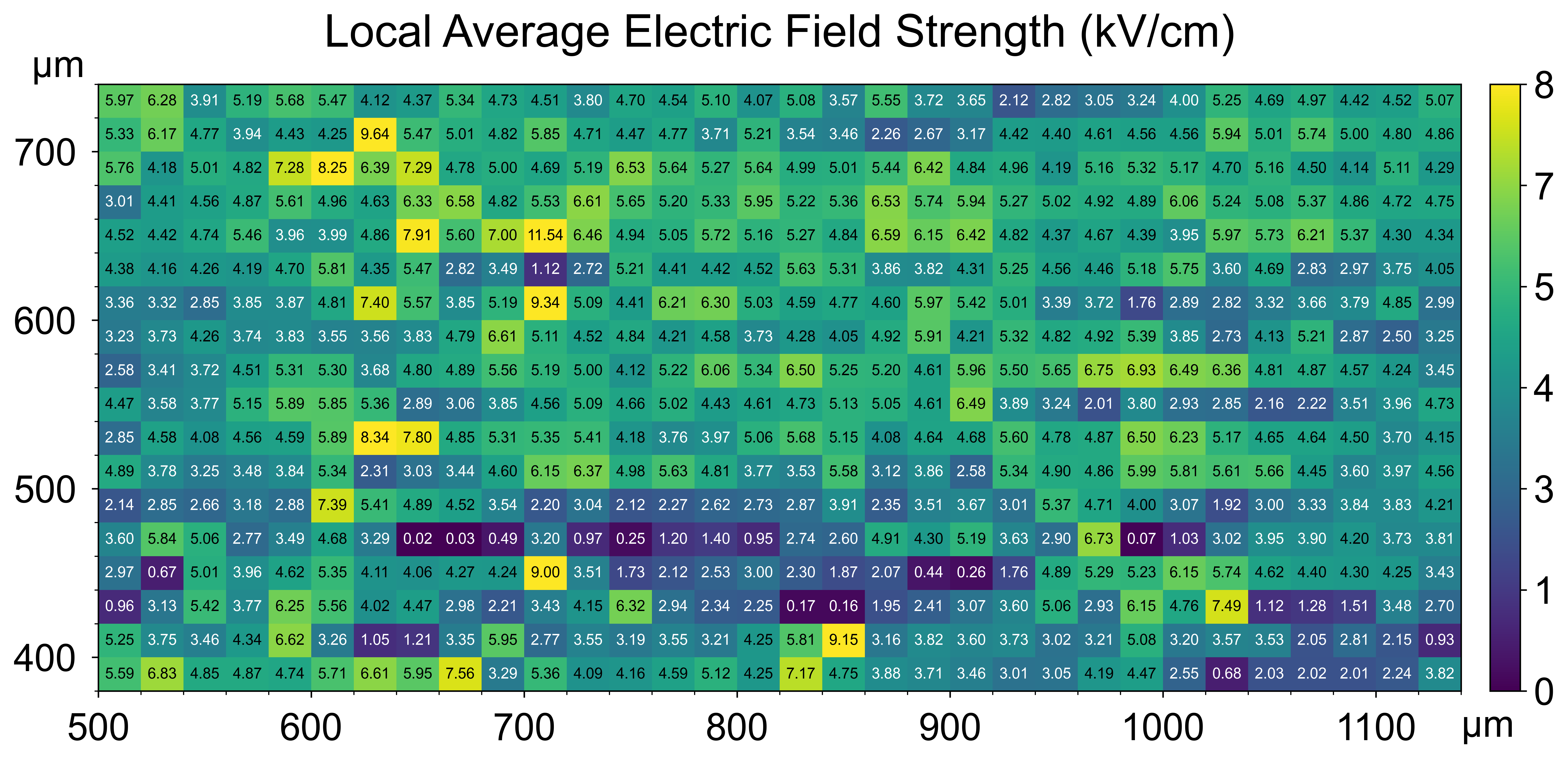
 **Figure S5** Spatial distribution of bin-averaged electric field magnitude. Field was averaged over 20 × 20 µm bins. Colored bins have an average field above the applied field, with color indicating magnitude.
